# Prototyping and Implementation of a Novel Feedforward Loop in a Cell-Free Transcription-Translation System and Cells

**DOI:** 10.1101/123190

**Authors:** Shaobin Guo, Richard M. Murray

**Affiliations:** Division of Biology and Biological Engineering, California Institute of Technology, Pasadena, California 91125, United States

## Abstract

Building novel synthetic biological devices is a time-consuming task because of the lengthy cell-based testing and optimization processes. Recent progress made in the cell-free field suggests that the utilization of mathematical models and cell-free transcription-translation testing platforms to systematically design and test novel synthetic biocircuits may help streamline some of the processes. Here we present a study of building a novel functional biological network motif from scratch with the aid of the mathematical modeling and the cell-free prototyping. In this work, we demonstrated that we were able to make a 3-promoter feedforward circuit from a concept to a working biocircuit in cells within a month. We started with performing simulations with a cell-free transcription-translation simulation toolbox. After verifying the feasibility of the circuit design, we used a fast assembling method to build the constructs and used the linear DNAs directly in the cell-free system for prototyping. After additional tests and assemblies, we implemented the circuit in plasmid forms in cells and showed that the *in vivo* results were consistent with the simulations and the outcomes in the cell-free platform. This study showed the usefulness of modeling and prototyping in building synthetic biocircuits and that we can use these tools to help streamline the process of circuit optimizations in future studies.

## Introduction

Traditional methods for building synthetic biological circuits are labor-intensive and time-consuming [1]. Recent research progress on a cell-free bimolecular breadboard platform provides us with a potential tool to perform fast circuit prototyping *in vitro* [2-4]. The *Escherichia* coli-based cell-free transcription-translation system (TX-TL) is a “biomolecular breadboard” that allows us to quickly design, build, test and debug novel synthetic biocircuits *in vitro* [5, 6]. Like a wind tunnel is to airplanes or a breadboard is to electronic circuits, TX-TL allows a biological engineer to quickly test, debug and retest their biological circuits *in vitro*, bypassing the time-consuming steps of cloning, transformation and cell growth, which are required for *in vivo* testing.

TX-TL is a cell-free system based on S30 cell extracts. The extracts have been optimized for *in vitro* biocircuits testing, which means it mimics the *E. coli in vivo* characteristics while preserving transcription, protein production capability and regulatory mechanisms [5, 6]. Previous work has shown that, besides plasmids, linear DNAs can also be used in TX-TL for fast circuit prototyping with the protection from the RecBCD inhibitor bacteriophage gamS protein [2, 7]. Combined this with the mathematical modeling, it is possible to rapidly design and characterize functional synthetic biocircuit modules, such as the gene regulation network motifs, in TX-TL.

Gene regulation networks are composed of a small set of recurring interaction patterns called network motifs [8, 9]. Among all the common gene regulation network motifs, the incoherent type-1 feedforward loop (FFL) is one of the widely used and interesting ones [10]. This specific FFL is composed of two input transcription factors, one (*x*) of which activates the other (*y*), and both jointly regulate (activate or repress) a target gene (*z*) (Figure 1A). Because there is a time delay between the activation of *z* by *x* and the repression of *z* by *y*, it has been shown that this type of FFL can generate a temporal pulse of *z* response.

**Figure 1:**
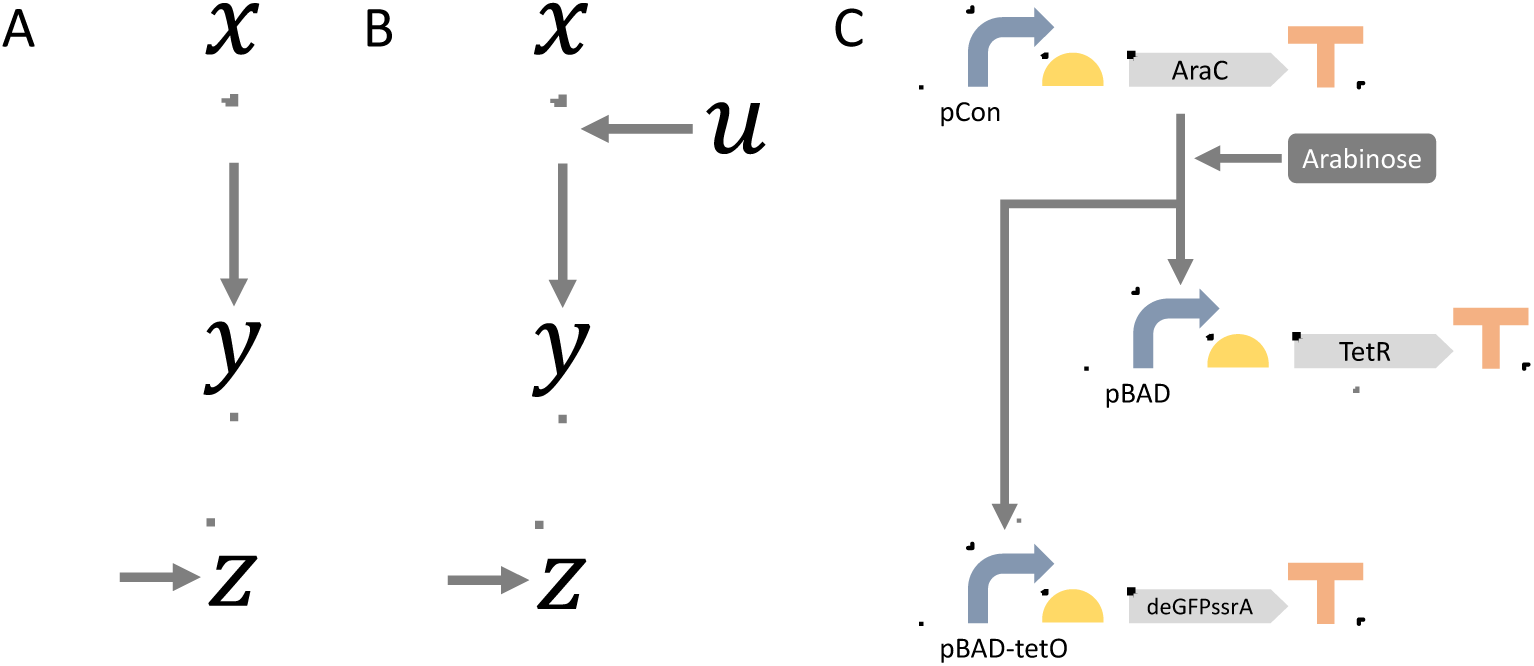
Diagrams of feedforward loops. **A:** The original feedforward loop illustration, composing of three components *x, y* and *z*. Arrows mean activation and bars mean repression. **B:** The adapted feedforward loop illustration, with additional input *u* to control the start of the activation. C: The actual biocircuit design for the feedforward loop. The activator AraC is at the 3’ of a constitutive promoter pCon. Arabinose is the inducer input. The repressor is controlled by the promoter pBAD, which is regulated by AraC/arabinose. The reporter deGFPssrA is at the 3’ of a combinatorial promoter pBAD-tetO, which can be activated by AraC/arabinose but can also be repressed by TetR.

Using this FFL as an example, here we demonstrate the process of using a mathematical model and the TX-TL prototyping platform to build novel synthetic biocircuits and characterize their behaviors. First, we verified our circuit design by performing simulations in a cell-free transcription-translation toolbox (TX-TL toolbox) [11]. Next, we built the constructs based on the verified design using a fast assembly method [12] and then we used the linear DNAs of these constructs directly for test in the TX-TL [2, 5]. After additional tests and assemblies, we implemented the FFL circuit *in vivo* and saw consistent results as *in silico* and *in vitro*. This study brought attention to utilizing mathematical models and the TX-TL prototyping platform when designing novel synthetic biocircuits. Instead of building and testing circuits directly in cells, we can save significant amounts of man-hours and streamline some of the prototyping steps involved in building new network motifs by properly employing this TX-TL system.

## Circuit design and simulations

To make things simpler in both TX-TL and subsequent cell-based tests, we first optimized the circuit design. Instead of having *x* directly turning on *y* and *z*, we added another component (*u*) to the circuit for better control (Figure 1B). For TX-TL and especially cell-based tests, we will put all three components of the circuit (*x, y* and *z*) into the testing platforms in the beginning. In order to control when the circuit should start the dynamics, an extra input is required. Without the presence of the input *u, x* cannot activate any of the downstream components. But as soon as *u* is added, the complex [*x*: *u*] will activate both *y* and *z* and initiate the dynamics. In this case, the activation of *y* and *z* is positively correlated with *u*, and *u* can be seen as an inducer that is required for *x* to activate the downstream parts.

Having the basic design in mind, we looked for biological parts that would fit the requirements stated above. We decided to use the AraC-arabinose activation system as [*x*:*u*] [13], a transcription factor TetR as the repressor *y* [14] and deGFP fluorescent protein with the corresponding combinatorial promoter as output *z* [15] (Figure 1C). The transcription factor AraC binds to the promoter pBAD and activates the transcription of downstream genes (TetR and deGFP) only in the presence of the inducer arabinose. On the other hand, the transcription factor TetR binds to operator site tet and represses the transcription of deGFP. A small molecule anhydrotetracycline (aTc) can be used to sequester TetR proteins away from the operator and resume the transcription. Because of the time delay between the activation by AraC-arabinose and the repression by TetR, deGFP gene will first be transcribed, and only after TetR proteins accumulate to certain threshold level (aTc can be used to extend the delay), the transcription of deGFP ceases. At the same time, the degradation ssrA tag on the deGFP protein leads to degradation of the protein [16]. As a result, the green fluorescence signal, which can be measured, will first increase and then decrease, generating a pulse-like behavior (Figure 2A).

**Figure 2:**
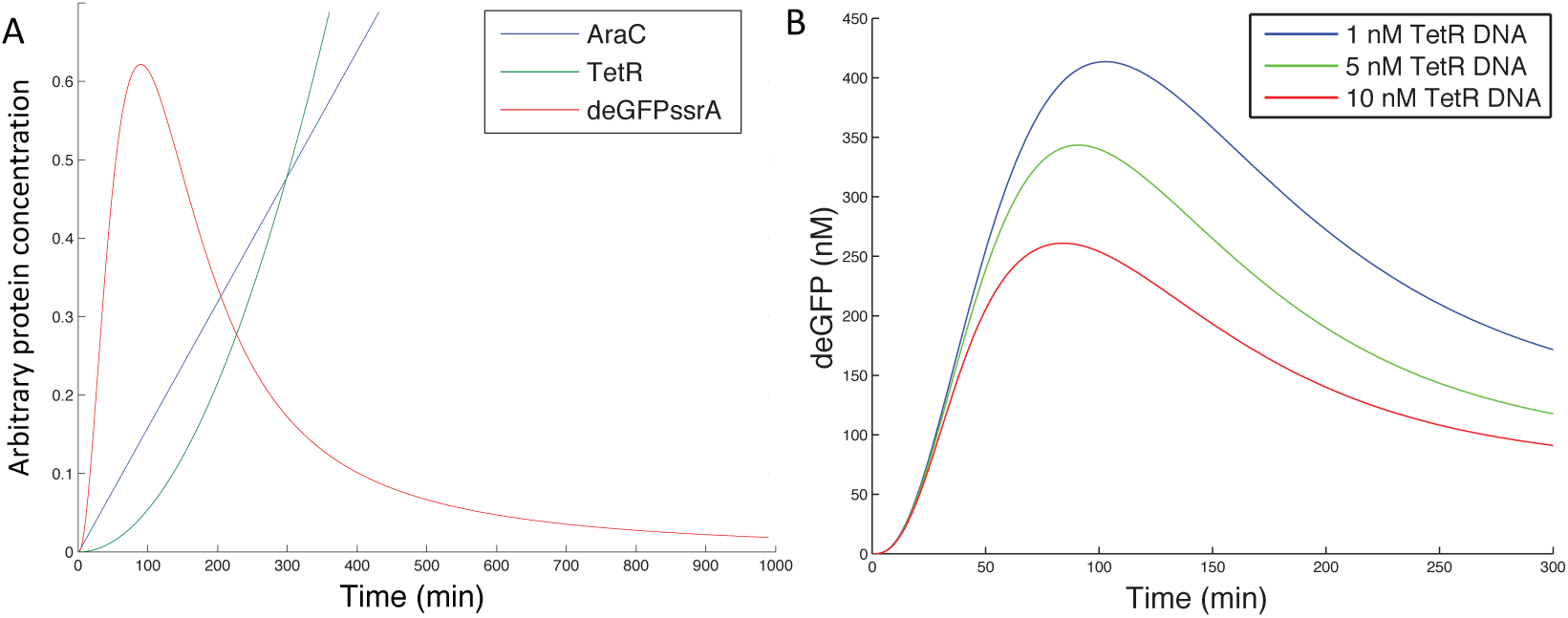
Simulation results of the FFL using the TX-TL simulation toolbox. **A:** Simulation of the time course of the arbitrary protein concentrations for all three circuit components. Initial AraC, TetR and deGFP DNA concentrations are set to be the same at 1. Arabinose concentration is also set to 1. AraC, which is controlled by a constitutive promoter, is assumed to be produced in a linear fashion. The pBAD promoter controlling deGFP is set to be stronger than the pBAD promoter controlling TetR, in order to extend the delay and produce significant amount of deGFP proteins. **B:** Simulation of the time course of the deGFP protein concentrations with varied initial TetR DNA concentrations from 1 nM to 10 nM. Initial conditions for AraC and deGFP were 10 nM and the arabinose concentration was set to be 0.27%.

Simulations were performed using the TX-TL toolbox [11]. By tuning the parameters in the toolbox model, we verified the feasibility of our design through simulation. In Figure 2A, we can see that the model captures the pulse-like behavior of deGFP protein concentration, along with the increasing activator AraC concentration and repressor TetR concentration. The delay in the increase of TetR concentration and the degradation of deGFP proteins result in the pulse-like behavior in deGFP concentration.

In both the mathematical model and the TX-TL platform, we have full control over the initial DNA concentrations for all the components. That gives us the freedom to change the circuit dynamics by simply changing the inputs – DNA concentrations. In Figure 2B, we tested how the varied concentrations of the repressor TetR DNA can affect the deGFP dynamics with all the other components as constants. As the simulation result shown here, increasing TetR DNA concentrations not only brings down the peak deGFP concentrations, but it also shifts the peak to the left, suggesting that it takes less time for TetR protein to accumulate to the threshold level when there is more initial TetR DNA.

## Linear DNAs and plasmids construction and prototyping in TX-TL

After verifying the circuit design using simulations, we decided to build the incoherent type-1 FFL shown in Figure 1C. We first designed and ordered primers (a day before) to amplify the coding sequences for AraC, TetR and deGFPssrA. Then we used the GoldenBraid assembly method to stitch specific promoters, ribosome binding sites (RBSs), coding sequences (CDSs) and terminators together with plasmid vectors [12]. After 1 hour incubation, we amplified the linear DNAs containing Promoter-RBS-CDS-Terminator-Vector via PCR reactions. Then we used these linear DNAs to run experiments in TX-TL directly with the presence of gamS. From start to finish, one experiment cycle can be less than a day.

TX-TL experiments were run by simply mixing the extract and buffer with the inputs – DNAs (AraC, TetR and deGFP) and the inducer arabinose [5]. After mixing all together, GFP fluorescence was measured using a plate reader. Figure 3A showed the experimental results from TX-TL experiments. All the curves were consistent with the simulation: GFP signal first increased as a result of AraC-arabinose activation; then after TetR proteins accumulated to the threshold amount, they repressed the transcription of deGFPssrA and at the same time, ClpX protein, which is an ATPase, unfolded the tagged deGFP proteins and caused the reduction of GFP signal [17]. As we can also see in the figure, the more the TetR DNAs were added, the lower the GFP signal was and the faster the signal reached peak; this was also consistent with our simulations. However, it is easy to notice that the steady state GFP protein concentrations for different initial TetR DNA concentrations are different. This is because TX-TL reactions have limited resources, including RNA polymerases, NTPs, ribosomes and amino acids [18]. The energy required for ClpX to unfold tagged deGFP proteins will run out gradually, and the resources will get used up faster when more DNAs are in the reaction. As a consequence, the GFP concentrations showed different steady state levels.

**Figure 3:**
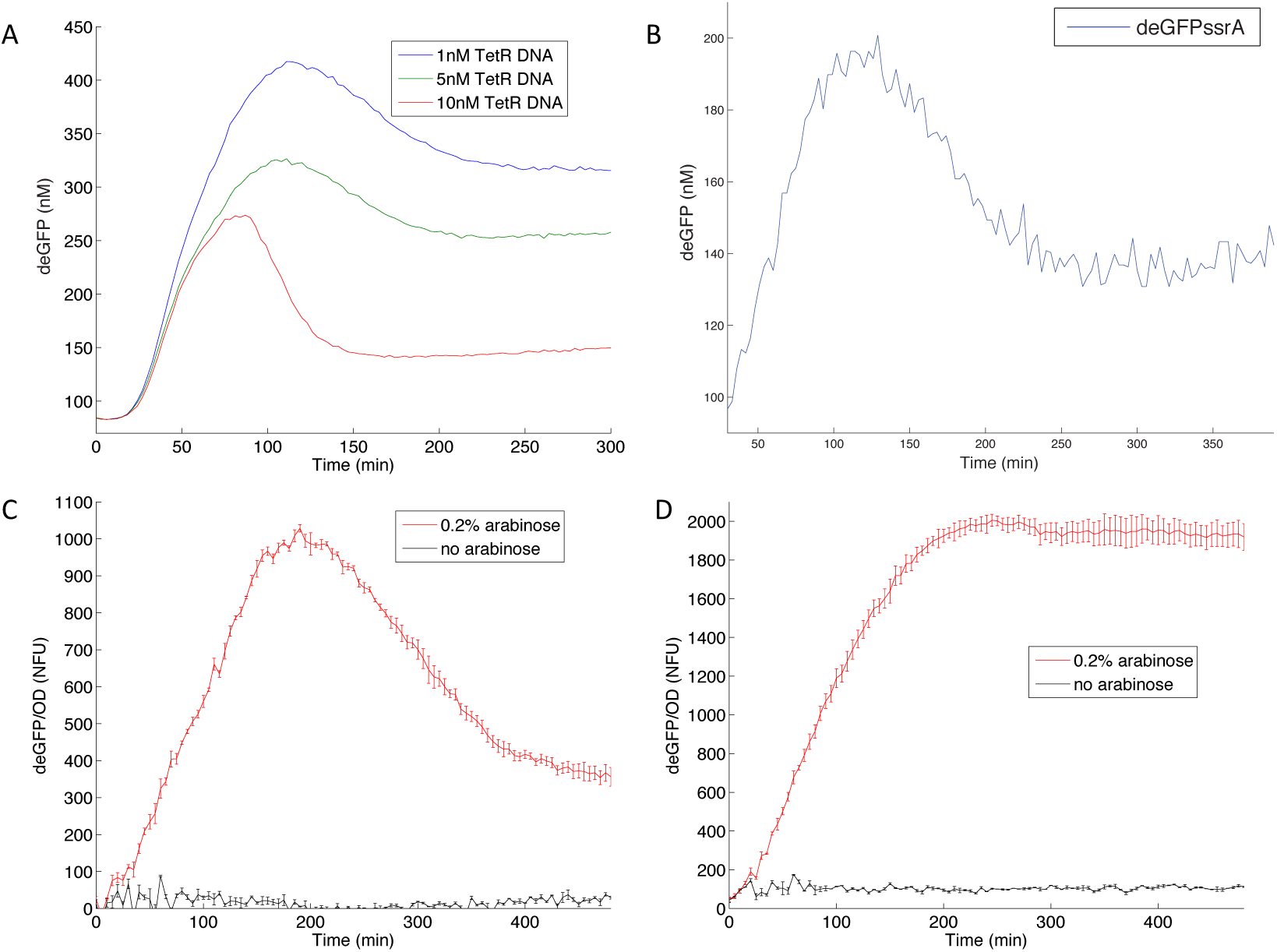
Experimental results of the FFL in TX-TL and cells. **A:** The time course result of the FFL experiment in TX-TL with linear DNA: 10 nM AraC linear DNA, 10 nM deGFP linear DNA, 10 nM ClpX linear DNA, 0.2% arabinose, 0.1 µg/mL aTc and varied TetR linear DNA concentrations. Experiments were run at 29°C. **B:** The time course result of the FFL experiment in TX-TL with plasmid DNA: 2 nM AraC-TetR-deGFPssrA plasmid, 1 unit of purified ClpX protein, 0.2% arabinose and 0.01 µg/mL aTc. Experiments were run at 29°C. **C, D:** The time course results of the FFL experiments in cells. **C:** deGFP protein is tagged with ssrA degradation tag. **D:** deGFP does not have a degradation tag. GFP signals, which were average from three independent repeated wells, were measured using a plate reader and then data were normalized using OD600 readings to get the fluorescence reading for each cell. The concentration of the inducer arabinose is 0 or 0.2%. Experiments were run at 37°C.

After we tested the linear DNA version of the FFL in TX-TL and found a working design, we assembled these linear DNAs into one plasmid using the GoldenBraid assembly method in order to implement the circuit in cells. Before testing the plasmid version of FFL *in vivo*, we first evaluated the construct in TX-TL, as it was fast and convenient to set up TX-TL experiments and there was no need for the time-consuming step of growing cells. Figure 3B showed the results of the plasmid version of the FFL in TX-TL. The dynamics of the circuit were again consistent with those from both simulations and linear DNA circuit, suggesting that this specific circuit design had a good chance to work in cells.

## Implementation of the biocircuit in cells

Following the test of expressing plasmid version of the FFL in TX-TL, we transformed that plasmid into *E. coli* cells. In addition, to make sure the decrease of GFP signal *in vivo* was specific to the GFP degradation by ClpX instead of the dilution introduced by cell division, we had a control circuit, in which the deGFP was not tagged with the ssrA degradation tag. Figure 3 showed the experimental results from *in vivo* experiments of the FFL circuit (Figure 3C) and the control circuit (Figure 3D). The dynamics shown in Figure 3C were clearly consistent with those shown in TX-TL experiments, meaning the FFL prototyped in TX-TL indeed showed the same behavior *in vivo*. In contrast, the control circuit in Figure 3D did not exhibit a pulse-like behavior, suggesting that the pulse we saw in Figure 3C was not a result of dilution but was caused by the degradation of the GFP proteins.

## Discussion

The idea of the cell-free TX-TL platform is not new as there have been many studies on the original S30 cell extract since it first came out in 1960s [19, 20]. However, most of the cell-free extracts were focusing on protein expression as an alternative option to making proteins in cells [21]. This specific TX-TL platform was developed with the goal of prototyping synthetic biocircuits in mind. TX-TL, along with the mathematical toolbox developed for it, can be the prototyping breadboard for synthetic biology. This work, together with other publications [2-4, 6, 18], serves as a testimony for that ambition. Expressing transcription factors and have them turning certain components on or off has been demonstrated before; but prototyping a synthetic biocircuit with spontaneous dynamics built in, such as an incoherent type-1 feedforward loop, is challenging. By tuning the parameters used by the mathematical model, we could get a sense of what strength of promoters and ribosome binding sites should be used in the actual biocircuit. Then we had to spend some amount of time characterizing specific parts, such as promoters, ribosome binding sites and coding sequences individually in TX-TL. Then from this preliminary parts library, we could assemble our constructs and test the actual components in TX-TL. Because of the fast iteration time of this platform, we were able to finish all the above and the final implementation in cells within a month. Compared to the first generation of biocircuits, we have shortened the development time significantly [22, 23]. We have also optimized a variant FFL circuit composed of a different activator in TX-TL and in cells (see “*Implementation of the feedforward loop composed of LasR and pLas”* in Supplementary Materials for details).

There are limitations to this protocol. First, there are limited resources in the TX-TL reactions and on top of that, there are competition of different components for the same transcription and translation machineries. One way to avoid this limitation is to set up reactions in compartments that only allow small molecules exchanges so that some resources, such as amino acids and NTPs, can be replenished. Second, it is challenging to use TX-TL data quantitatively to deduce the results in cells. Cell-free systems, no matter how we justify it, are different from cells. When TX-TL is used to help with prototyping novel synthetic biocircuits, it is recommended that the results are examined qualitatively instead of quantitatively. Although the absolute strengths of certain promoters and RBSs are different between TX-TL and cells, their relative strengths are comparable between the two systems. Third, the TX-TL simulation toolbox has its uncertainty and arbitrariness. The parameters used in the toolbox might not be physiologically reasonable despite the fact that we tried to refer to as many literature available parameters as possible. However, qualitatively we can use the simulation results as a reference, for example, we need one promoter to be stronger than the other promoter in order to achieve the desired dynamics and then we can have this information in mind when we design the actual biocircuits. In summary, though it is not a platform without its limitations, TX-TL could certainly be used for rapid preliminary characterization and prototyping of synthetic biocircuits.

## Materials and Methods

### Cell-free experiment preparation and execution

Preparation of the cell-free TX-TL expression system was done according to previously described protocols [5], resulting in extract with conditions: 8.9 - 9.9 mg/mL protein, 4.5 - 10.5 mM Mg-glutamate, 40 - 160 mM K-glutamate, 0.33 - 3.33 mM DTT, 1.5 mM each amino acid except leucine, 1.25 mM leucine, 50 mM HEPES, 1.5 mM ATP and GTP, 0.9 mM CTP and UTP, 0.2 mg/mL tRNA, 0.26 mM CoA, 0.33 mM NAD, 0.75 mM cAMP, 0.068 mM folinic acid, 1 mM spermidine, 30 mM 3-PGA, 2% PEG-8000.

TX-TL reactions were conducted in a volume of 10 µL in a 384-well plate (Nunc MicroWell 384-well optical bottom plates) at 29°C, using a three-tube system: extract, buffer, and DNA. When possible, inducers such as arabinose or purified proteins such as gamS [7] were added to a mix of extract and buffer to ensure uniform distribution. For deGFP, samples were read in a Synergy H1 plate reader (Biotek) using settings for excitation/emission: 485 nm/525 nm, gain 61 or 100. All samples were read in the same plate reader, and for deGFP relative fluorescent units were converted to either nM (for TX-TL) or Normalized Fluorescent Unit (NFU for *in vivo*) using a purified deGFP-His6 standard to eliminate machine to machine variation (different Bioteks).

### PCR product preparation and plasmid DNA assembly

Linear DNA fragments were amplified using Pfu Phusion Polymerase (New England Biolabs), DpnI digested for 5 min at 37°C (New England Biolabs) while verified with agarose gel electrophoresis, and PCR purified using previously described procedures. Fragments were then assembled *in vitro* using Golden Gate assembly. For Golden Gate assembly, a 15 µL reaction was set up consisting of equimolar amounts of vector and insert, 1.5 µL 10X NEB T4 Buffer (New England Biolabs), 1.5 µL 10X BSA (New England Biolabs), 1 µL BsaI (New England Biolabs), and 1 µL T4 Ligase at 2 million units/mL (New England Biolabs). Reactions were run in a thermocycler at 10 cycles of 2 min/37°C, 3 min/20°C, 1 cycle 5 min/50°C, 5 min/80°C. For Golden Gate assembly, constructs with internal BsaI cut sites were silently mutated beforehand using a QuikChange Lightning Multi Site-Directed Mutagenesis kit (Agilent). For both methods, assembled circular DNAs were transformed into electrocompetent or chemically competent cells: a KL740 strain (lab made competent strain) if using an OR2-OR1 promoter (29°C), a MG1655 strain (lab made competent cells) for circuit testing, and a JM109 strain (Zymo Research) for all other constructs. KL740 upregulates a temperature sensitive lambda cI repressor. PCR products were amplified using Pfu Phusion Polymerase (New England Biolabs) for all constructs, and were DpnI digested. Plasmids were miniprepped using a Qiagen mini prep kit. All plasmids were processed at stationery phase. Before use in the cell-free reaction, both plasmids and PCR products underwent an additional PCR purification step using a QiaQuick column (Qiagen), which removed excess salt detrimental to TX-TL, and were eluted and stored in 10 mM Tris-Cl solution, pH 8.5 at 4°C for short-term storage and −20°C for long-term storage. All the plasmids used in the work can be found on https://www.addgene.org/.

### *In vivo* experiment

All *in vivo* experiments were performed in *E. coli* strain MG1655. Plasmid combinations were transformed into chemically competent *E. coli* MG1655 cells, plated on Difco LB+Agar plates containing 100 µg/mL carbenicillin and incubated overnight at 37°C. Plates were taken out of the incubator and three colonies were picked and separately inoculated into 5 mL of LB containing carbenicillin and/or chloramphenicol, and/or kanamycin at the concentrations above in a 14 mL Falcon Round-Bottom Polypropylene Tubes (Fisher Scientific), and grown approximately 17 h overnight at 37°C at 200 rpm in a benchtop shaker. This overnight culture (100 µL) was then added to a new 14mL tube containing 5 mL (1: 50 dilution) of Minimal M9 casamino acid (M9CA) media [1X M9 salts (42 mM Na_2_HPO_4_, 24 mM KH_2_PO_4_, 9 mM NaCl, 19 mM NH_4_Cl, 1 mM MgSO_4_, 0.1 mM CaCl_2_, 0.5 µg/ml thiamine, 0.1% casamino acids, 0.4% glycerol) containing the selective antibiotics and grown for 4 h at the same conditions as the overnight culture. Then 10 µL cultures were transferred to 96-well glass bottom plate with 290 µL M9 with corresponding experimental conditions. Plates were shaken and GFP fluorescence (485 nm excitation, 525 nm emission), and optical density (OD, 600 nm) were measured using a Biotek Synergy H1m plate reader at 37°C at the highest speed for 12 hours.

## Funding Sources

This work was supported by DARPA Living Foundries (Grant number: HR0011-12-C-0065).

## Notes

R.M.M. has ownership in a company that commercializes the cell-free technology utilized in this paper. All the other authors claim no competing interest.

## ACKNOWLEDGMENT

We would like to thank Clarmyra Hayes, Yutaka Hori, Vipul Singhal for helpful discussion and suggestions. We would like to thank Murray lab members for useful suggestions.

## Supplementary Materials

### Implementation of the feedforward loop composed of LasR and pLas

Besides the feedforward loop (FFL) circuit we designed, prototyped and implemented in the main article, we also tested a FFL designed by Zachary Sun, which had AraC, arabinose, pBAD and pBAD-tetO replaced by LasR, N-(3-Oxododecanoyl)-L-homoserine lactone (AHL), pLas and pLas-tetO, respectively. The circuit design is shown in Supplementary Figure S1.

**Supplementary Figure S1.**
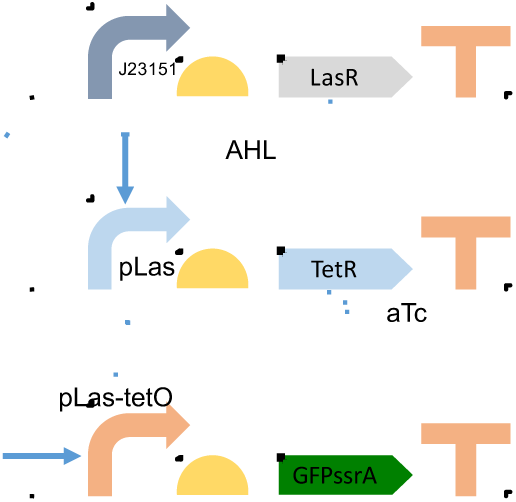
Illustration diagram of the FFL composed of LasR. The arrows mean activation and the bars mean repression.

The promoter J23151 is a constitutive promoter [24]. LasR protein, in the presence of the inducer AHL, becomes an activator that can turn on both pLas and pLas-tetO promoters. All the other components work like the AraC FFL and the time course data of the LasR circuit from *in vivo* experiments are shown in Supplementary Figure S2.

**Supplementary Figure S2.**
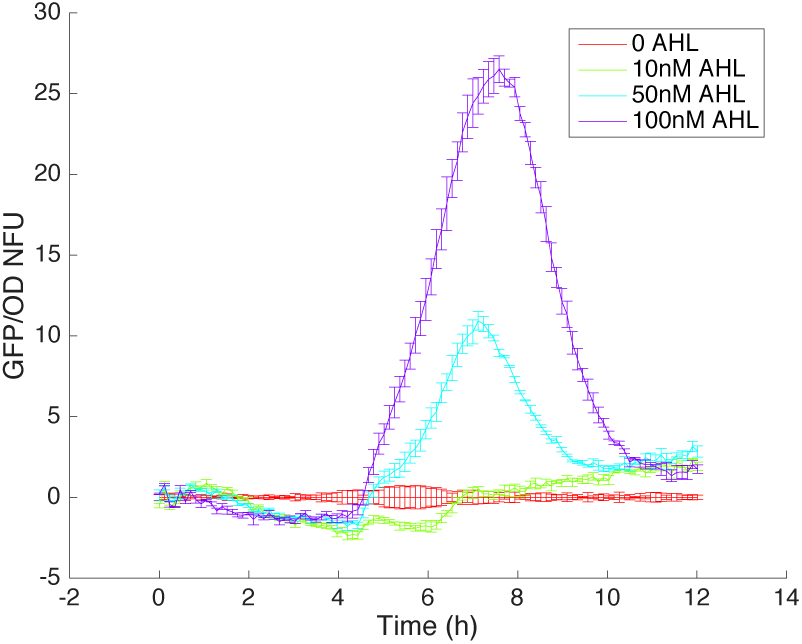
Time course data from the *in vivo* experiment of the LasR FFL circuit in MG1655 *E. coli* cells. Detailed experiment setup was described in the Materials and Methods. Briefly, 4 AHL concentrations were used in the experiment, from 0 to 100 nM. 20 ng/mL of aTc was added to all of them. GFP fluorescence and OD600 measurements were done using a Biotek plate reader. GFP data was subtracted by the background and then normalized with OD data to get the normalized fluorescence unit (NFU).

Though the LasR FFL works in a similar way as the AraC FFL, one problem with the LasR FFL circuit was that the aTc, which would bind to TetR proteins and sequester them away from tetO, had to be added to extend the delay to create significant pulses. This was due to the leaky expression of the pLas promoter. To make the circuit more robust, we engineered the pLas promoter to be more tightly controlled by LasR-AHL via prototyping different variants of pLas promoters in TX-TL. The one we found working very robustly is shown in Supplementary Figure S3 and the sequence of that pLas variant can be found on https://www.addgene.org/. As we can see in the figure, only when both LasR and AHL were added, we could see the activation of the pLas-GFP. Neither LasR nor AHL alone could activate the promoter and there was little to none leaky expression from the promoter itself. TetR and aTc were tested to make sure that the pLas promoter, without the tetO part, could not be affected by them.

**Supplementary Figure S3.**
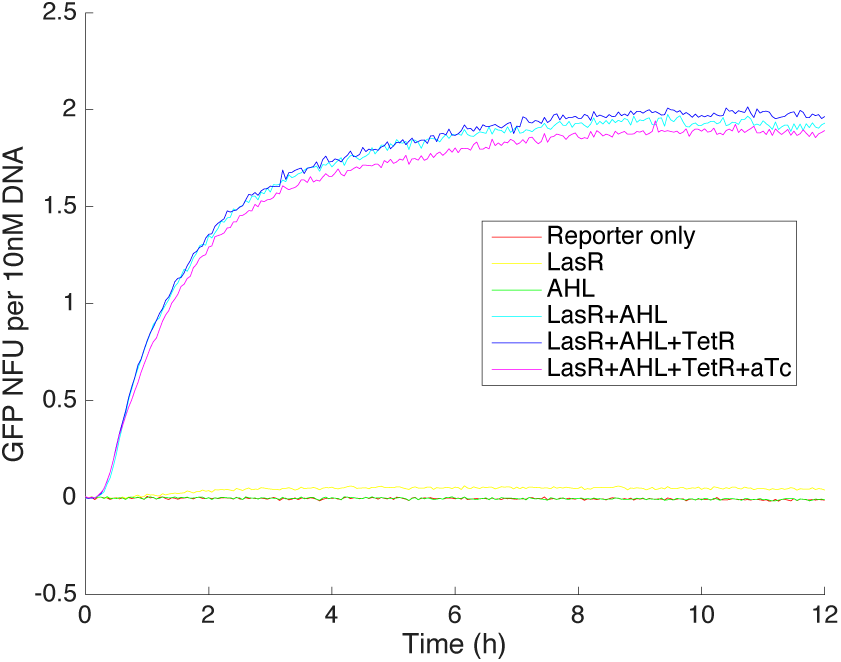
Time course of the selected pLas promoter with GFP at its 3’. Experiments were run with 10 nM pLas-GFP linear DNA with or without the additional components/inducers listed in the legend. If used, the pCon-LasR linear DNA was 10 nM, pCon-TetR linear DNA was 10 nM, AHL was 50 nM and aTc was 20 ng/mL. The GFP fluorescence data was subtracted with background and then normalized using GFP protein calibration data on a Biotek plate reader.

We then modified the LasR FFL circuit with this exact promoter and we were able to generate pulse-like behavior in cells without adding any aTc. The *in vivo* data is shown in Supplementary Figure S4. As we can see, when there was no AHL added, we got no response from the circuit. Only when we had significant activation caused by AHL (more than 50 nM in this case), we could see pulse-like behavior from the circuit in cells. And the highest peaks of the pulses were positively correlated with the inducer concentrations.

**Supplementary Figure S4.**
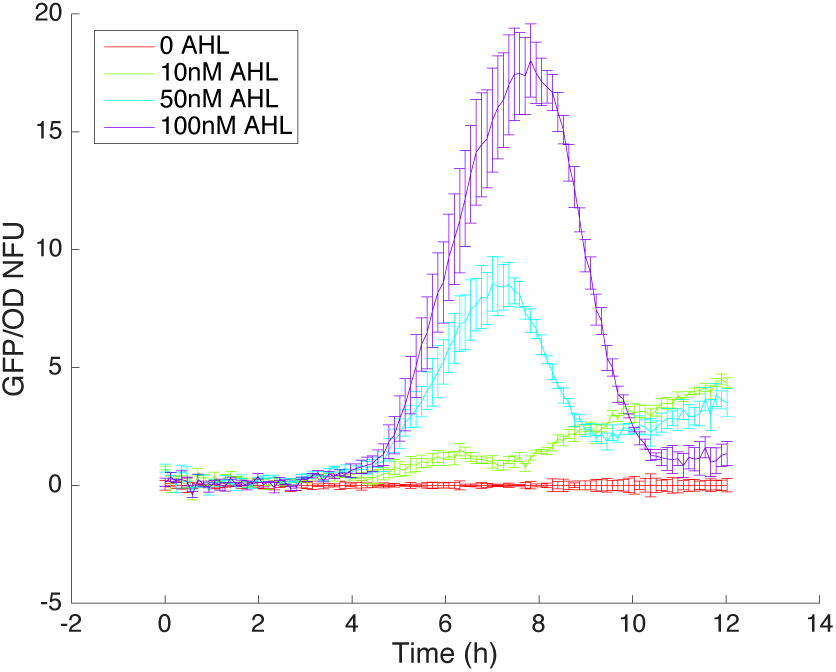
Time course data from the *in vivo* experiment of the optimized LasR FFL circuit in MG1655 *E. coli* cells. Detailed experiment setup was described in the Materials and Methods. Briefly, 4 AHL concentrations were used in the experiment, from 0 to 100 nM. 20 ng/mL of aTc was added to all of them. GFP fluorescence and OD600 measurements were done using a Biotek plate reader. GFP data was subtracted by the background and then normalized with OD data to get the normalized fluorescence unit (NFU).

Not only did we test the circuit quantitatively in bulks, but we also examined the circuit behavior qualitatively in a microfluidic device, also known as the mother machine. The mother machine consists of a series of growth channels that can trap single bacterial cells inside, and is designed to allow growth medium to pass through at a constant rate, which results in diffusion of fresh medium into the growth channels as well as removal of cells as they emerge from the channels into the main trench [25]. As we can see in Supplementary Figure S5, while cells growing and dividing in the narrow comb-like channel, they showed green fluorescence intensity starting from weak to strong and then back to weak with the same media keeping the inducer AHL concentration constant at 50 nM.

**Supplementary Figure S5.**
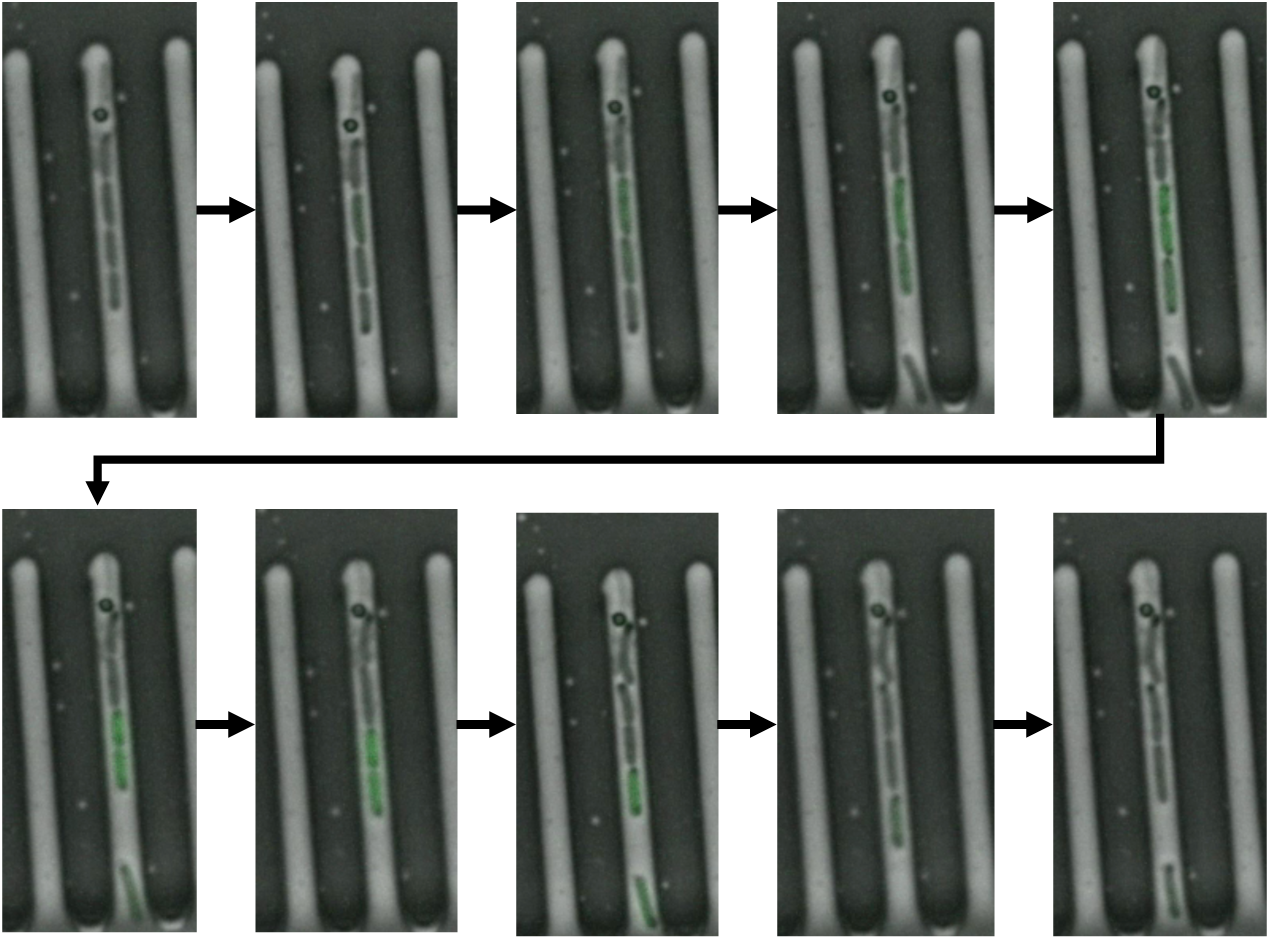
Microscope movie snapshots of the optimized LasR FFL. Each frame was taken 10 minutes apart. Cells were grown using the same *in vivo* method described in the Materials and Methods. Fluorescence microscopy imaging was performed on an Olympus IX81 inverted fluorescence microscope using a Chroma wtGFP filter cube (450/50 BP excitation filter, 480 LP dichroic beamsplitter, and 510/50 BP emission filter), with an XFO-citep 120 PC light source at 100 % intensity and a Hamamatsu ORCA-03G camera. Cells were imaged using a 100x phase objective with oil.

